# Electrophysiological Brain Network Estimation with Simultaneous Scalp EEG and Intracranial EEG: Inference Algorithm and Applications

**DOI:** 10.1101/2024.04.16.589846

**Authors:** Shihao Yang, Feng Liu

## Abstract

Activity in the human brain is composed of complex firing patterns and interactions among neurons and neuronal circuits. The neuroimaging field underwent a paradigm shift over the past decades from mapping tasked evoked brain regions of activations towards identifying and characterizing the dynamic brain networks of coordinating brain regions. Electrophysiological signals are the direct manifestation of brain activities, thus characterizing the whole brain electrophysiological networks (WBEN) can serve as a fundamental tool for neuroscience studies and clinical applications. The electrophysiological network inferred from electroencephalogram (EEG) source imaging suffers from low accuracy limited by the Restricted Isometry Property (RIP), while the invasive EEG-derived electrophysiological networks can only characterize partial brain regions where invasive electrodes reside. In this work, we introduce the first framework for the integration of scalp EEG and intracranial EEG (iEEG) for WBEN estimation with a principled estimation framework based on state-space models, where an Expectation-Maximization (EM) algorithm is designed to infer the state variables and brain connectivity simultaneously. We validated the proposed method on synthetic data, and the results revealed improved performance compared to traditional two-step methods using scalp EEG only, which demonstrates the importance of the inclusion of iEEG signal for WBEN estimation. For real data with simultaneous EEG and iEEG, we applied the developed framework to understand the information flows of the encoding and maintenance phases during the working memory task. The information flows between the subcortical and cortical regions are delineated, which highlights more significant information flows from cortical to subcortical regions compared to maintenance phases. The results are consistent with previous research findings, however with the view of the whole brain scope, which underscores the unique utility of the proposed framework.

## 1 Introduction

Brain connectivity networks consist of complex connections between neurons that carry information throughout the brain and support various cognitive functions in neuroscience [1, 2]. However, estimating and reconstructing these connectomes is a complex task that requires a combination of multiple neuroimaging and computational modeling techniques [3–6]. Many studies have suggested that an accurately inferred brain connectivity map can gain insights into the relationship between brain structure and function, reveal the fundamentals of cognitive processes, and uncover the mechanisms of neuropsychiatric diseases [7–12].

In the past decades, most researchers leveraged functional Magnetic Resonance Imaging (fMRI) for brain functional network modeling and analysis. fMRI is one of the most widely used non-invasive neuroimaging techniques, it indirectly measures the brain activity by measuring changes in blood flow dynamics as well as oxygen levels. fMRI can provide a high spatial resolution of activity. fMRI has been used to characterize the functional connections between different regions of the brain [13–15]. By analyzing fMRI data, researchers can explore the connectomes of the brain under different states and tasks, unveiling the complex relationships between the structure and function of brain networks [15–17]. However, fMRI has limitations on relatively high cost, non-portability, and low temporal resolution, which limit its applications where high temporal solution is necessary [18, 19]. Therefore, the neuroimaging techniques with a temporal resolution of subsecond level, such as Magnetoencephalography (MEG) or Electroencephalogram (EEG) can be used to estimate the WBEN after Electrophysiological Source Imaging (ESI). Studies that use MEG or EEG to reconstruct brain functional networks have leveraged different ESI approaches and connectivity metrics [20–26]. Those frameworks consists of two steps, which is Step (1): EEG/MEG source localization or ESI; Step (2): brain network construction with the estimated source space signal.

The existing ESI framework suffers from low accuracy when it comes to the estimation of whole brain regions due to the ill-posed property and is limited by Restricted Isometry Property (RIP) in theory [27]. Traditionally, neurophysiologically plausible priors, such as sparsity, have to be incorporated in ESI as the regularization terms to enforce a feasible solution [28–30]. Estimation of the whole brain networks poses more challenges to solving the ESI problem as it further relaxes the sparsity constraint [31]. Another existing problem for the current ESI framework is there is no supervision in the learning process of the latent source space signal, and model validation is conducted only after the reconstruction of the source signal based on either synthetic data or neuroscience knowledge, thus limiting the usability and accuracy of the estimated source signal.

In addition to non-invasive modalities, invasive neuroimaging technologies, such as intracranial Electroencephalography (iEEG) including Electrocorticography (ECoG) and stereoelectroencephalography (sEEG) that place subdural electrodes on the brain surface or penetrating electrodes in the subcortical brain regions can achieve a more accurate connectivity map among the regions of interest within the brain of high temporal resolution, in order to mitigate the noise caused by extracranial measurements [32–34]. However, it is infeasible to record the whole brain’s intracranial signal due to the invasive nature of iEEG, which makes the brain only partially observable. As iEEG electrodes can only cover part of the brain, important nodes or links might be missing at other brain regions. For example, in the epilepsy seizure onset zone analysis, Gao et al. used MEG/EEG source imaging and identified the interictal spikes that are missed by the iEEG [35], which showcases the value of a synergy of scalp EEG and invasive iEEG. Another group of researchers suggests most interictal epileptiform discharges arise in brain tissue outside the site of seizure onset and propagate toward it [36]. In such cases, having a WBEN estimation can provide a more comprehensive characterization of signal genesis and propagation at the whole brain scale.

To address the above issues, we take the scalp EEG and iEEG as complementary to each other and believe the fusion of two modalities can provide a delineation of the whole brain’s activation/connectivity map[35, 37]. The integration of iEEG and EEG can yield a faithful reconstruction of whole brain electrophysiological connectivity, however, a principled multimodal integration modeling and inference framework has not been explored before. The pipeline of the proposed approach is illustrated in Fig. 1

**Fig. 1.**
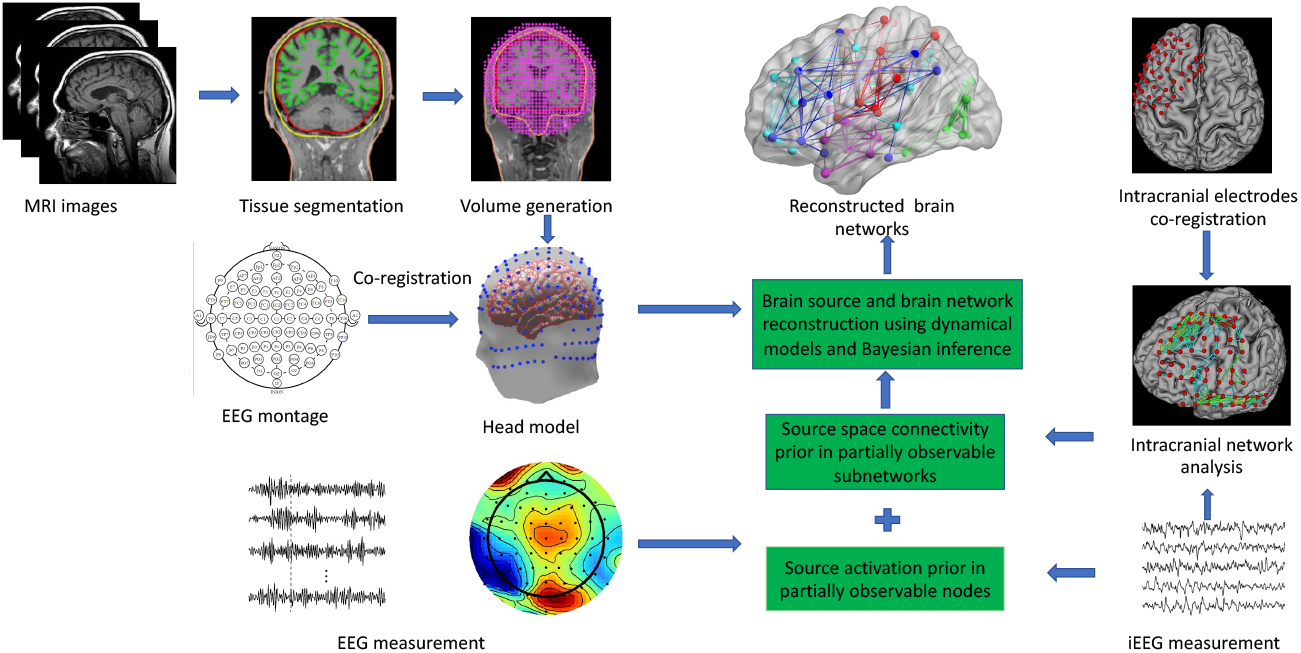
The overall pipeline of integration of EEG and iEEG for source space brain network reconstruction

## 2 Method

### 2.1 Basic problem definition

The linear discrete dynamic system of source as well as the linear model of EEG and iEEG observations can be defined as:

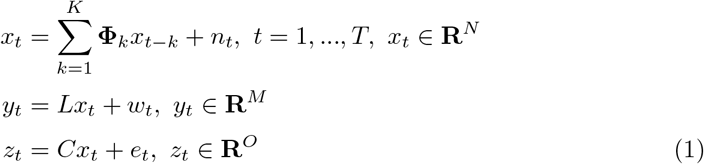

where *N, M*, and *O* are the number of the source regions, EEG electrodes, and iEEG electrodes, respectively. **Φ**_*k*_ ∈ **R**^*N×N*^ is the state transition matrix that delineates the impact of the source state at time *t − k* to *t. n*_*t*_ ∈ **R**^*N*^, *n*_*t*_ *∼ 𝒩* (**0, Q**) is the noise in source state space which is assumed to be a multivariate Gaussian distribution with mean 0 and diagonal covariance matrix **Q**. *L* ∈ **R**^*M×N*^ is the lead field matrix. *w*_*t*_ ∈ **R**^*M*^, *w*_*t*_ *∼ 𝒩* (**0, P**) is the measurement noise in EEG observation which is also assumed to be a multivariate Gaussian distribution with mean 0 and a covariance matrix **P** that is assumed to be known by measuring on a realistic head model. And *C* ∈ **R**^*O×N*^ is a full-row rank transformation matrix that selects the source signal where its region can be observed by iEEG directly. *e*_*t*_ ∈ **R**^*O*^, *e*_*t*_ *∼ 𝒩* (**0, S**) is the iEEG observational noise which is assumed to follow multivariate Gaussian distribution with mean 0 and a covariance matrix **S** that is also can be measured in a similar manner as **P**. Thus, we can see from the model definition that the parameters that need to be estimated are **Φ** and **Q**.

Define the unknown parameters as *θ* = {**Φ, Q**}. Then the log-likelihood can be written in the form:

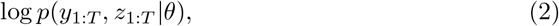

The log-likelihood of the model can be defined as

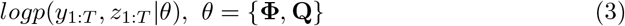

where **Φ** and **Q** are the parameters for the model, and its definition is the same as before.

Since the number of observations is far more than that of the source, the inverse estimation problem is highly ill-posed. To alleviate this phenomenon and simplify the problem, we add a regularization term based on the assumption that the connection from all other regions to a given region is sparse. In this case, we can define the regularized maximum log-likelihood of parameters for the model as :

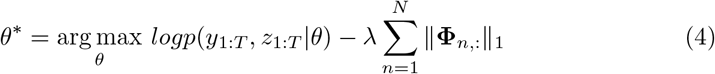

where Φ_*n*,:_ is the *n*^*th*^ row of the state transition matrix. *λ* is a regularization weight for model estimation that can be decided manually according to the experience or by grid search [38].

### 2.2 An EM estimation framework

In the domain of statistical methodology, the Expectation–Maximization algorithm serves as an iterative computational approach meticulously employed for the purpose of ascertaining the utmost likelihood, whether it pertains to local optima or the acme of posterior estimations (MAP) of parameters within the context of statistical models. These models, notably contingent upon latent variables concealed from direct observation, underscore the necessity for such intricate methodologies. The EM iterative procedure is characterized by its cyclical execution, entailing two pivotal stages: firstly, the Expectation (E) step, whereby a function is crafted to encapsulate the expected value of the log-likelihood, thoughtfully computed utilizing the prevailing parameter estimate. Secondly, the Maximization (M) step, during which a judicious parameterization is derived, endeavoring to maximize the anticipated log-likelihood cultivated during the antecedent E step. Subsequently, these diligently derived parameter estimates assume a central role in illuminating the distributional characteristics of the latent variables in preparation for the ensuing E step within this iterative loop. Since the data distribution in the source domain is unknown, we can apply the Expectation-Maximization(EM) framework that takes the source state as the latent variable to find the optimal estimation of the log-likelihood and then obtain the approximated state in the source domain with the problem defined above.

We will firstly illustrate the E-step, by starting from the Equation 2 defined above and using the facts that the observations of EEG and iEEG are conditional independent on the source state *x*, we can rewrite and derive the log-likelihood by introducing source state *x*_1:*T*_ as

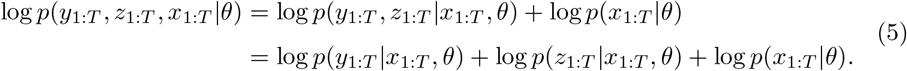

The first and second terms of the right-hand side of the second row can be easily obtained based on the Gaussian assumption of the noise while source *x*_*t*_ is given. Thus we can get

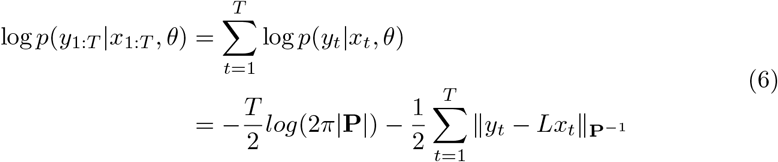

and

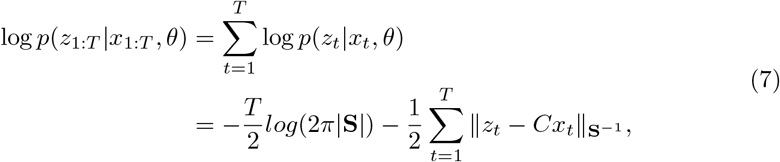

where | *·* | is the matrix determinant, and ∥*v*∥_*W*_ = *v*^*T*^ *Wv* is the quadratic form in the exponential term of multivariate Gaussian distribution. The last term can also be obtained in a similar way based on the linear dynamical model from Equation 1. and utilizes the presumption that **Q** is a diagonal covariance matrix with the items 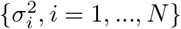 on its diagonal, then we have

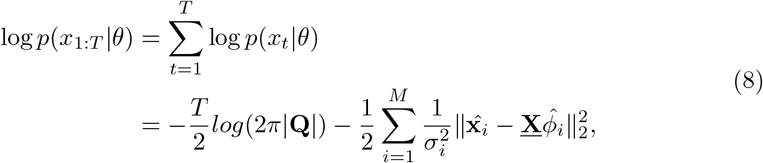

where 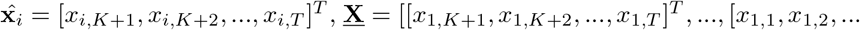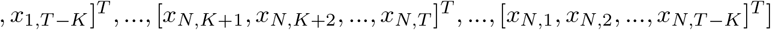 and 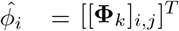 for *k* = 1,…,*K, j* = 1,…,*N*. By substituting Equations 6-8 into 5, we can reformulate the formula and take the expectation to get the Q-function for EM as

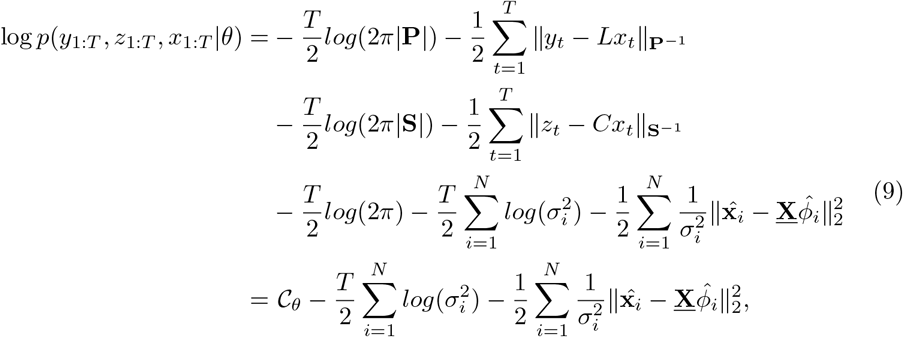

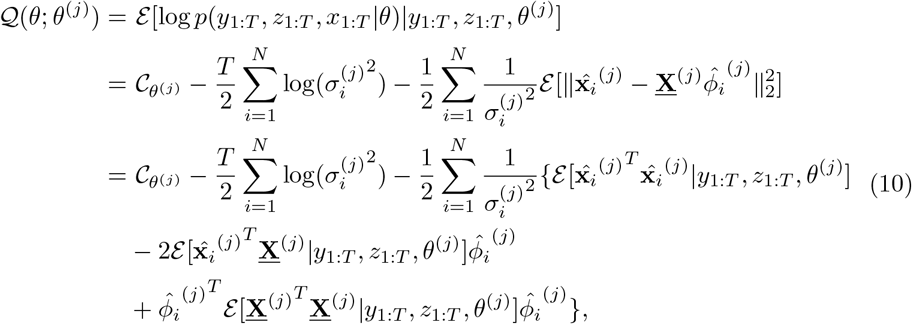

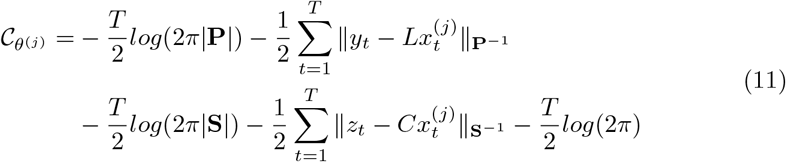

where the bracket superscript represents the *j*^*th*^ iteration in EM, and *C*_*θ*(*j*)_ is a constant term when *θ* is given at *j*^*th*^ iteration. We can find that the *p*(*x*_1:*T*_ |*y*_1:*T*_, *z*_1:*T*_, *θ*) is also Gaussian due to the Gaussian property on *x, y*, and *z* given *θ* [39]. In order to find the Q-function, we can permute the equation and notice that the first and third expectation terms in the last row of the Equation 10 consist of the second-order moment of the density *p*(*x*_1:*T*_ |*y*_1:*T*_, *z*_1:*T*_, *θ*) while the second expectation term can be expressed by the first-order moment of *p*(*x*_1:*T*_ |*y*_1:*T*_, *z*_1:*T*_, *θ*) whose mean as well as the covariance matrix can be estimated via Fixed Interval Smoothing (FIS), and the details of it will be listed in the next section.

In the M-step, we need to find the optimal *θ* = {**Φ, Q**} that maximizes the Q-function defined above. Since the number of observations is far less than that of the source regions which makes the inverse problem ill-posed. We then introduce the *l*_1_ regularization on **Φ** based on the premise that the functional connection among a given region and others possesses sparse properties to reduce the difficulty of problem-solving. Thus, we can have the equation of the maximization with the form

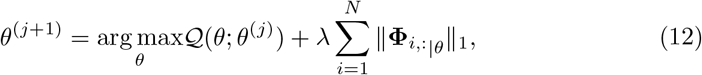

which can be solved easily by using the Fast Adaptive Shrinkage/Thresholding Algorithm (FASTA). In this way, we successfully build up the EM framework to estimate the source.

### 2.3 FIS for source density estimation

Fixed interval smoothing is a statistical technique used in time series analysis and signal processing. It involves retrospectively estimating and improving the values of a time series over fixed time intervals, considering both past and future observations.

This method is particularly useful for reducing noise or uncertainty in historical data and obtaining more accurate, smoothed estimates of the underlying trends or states within the time series.

The principle of FIS consists of two parts, forward filtering and backward smoothing. The forward filter is executed to derive posterior estimates and covariances up to the given time *t*. Subsequently, the backward filter is applied to yield prior estimates and covariances, effectively extending the timeline backward to time *t* or providing a prior perspective in reverse chronology, in other words. Finally, the estimates and covariances derived from both forward and backward filtering at time *t* are integrated to produce the ultimate estimation of the state and covariance matrix.

Recall the main problem in Equation 1. Taking into account computing performance factors, we can define a fusion observation by combining *y* and *z* together. The intuition is that we can think of EEG and iEEG as one measurement, but the measuring noise between two types of electrodes has no correlation, and the conditional density of merged observation is also a Gaussian. Thus, we redefine observations as

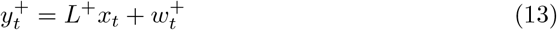

where

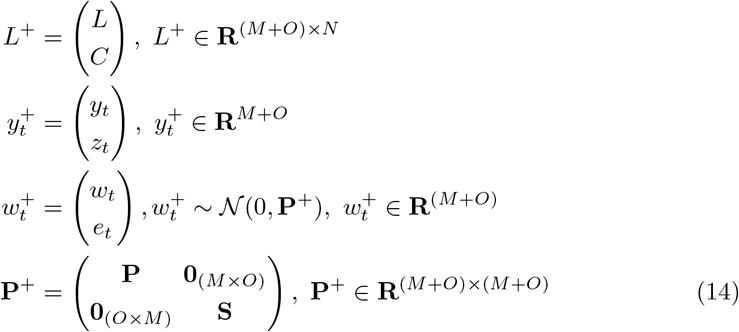

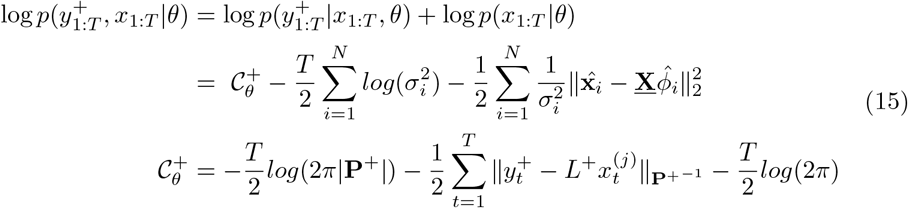

Then the log-likelihood problem can be transformed to Equation 15 and the same transformation can be applied to Q-function. The estimation of the mean and the covariance matrix of *p*(*x*_1:*T*_ |*y*_1:*T*_, *z*_1:_ *θ*) can be redefined as that of the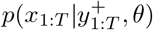. And we can easily find that 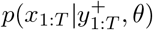. is a Gaussian, since for two jointly Gaussians, the conditional distribution is also a Gaussian [39]. Then, we can apply the FIS to find the mean and covariance matrix for 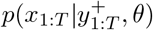. The forward Kalman filtering can give the estimation on 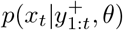 ∀*t*, while the backward Kalman smoothing can calculate the 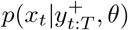 ∀*t*. The merging of two estimates can generate the final estimate on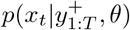 ∀*t*. Next, referring to the estimation framework[38], we start with Vector Auto Regressor (VAR) to generate the initial value for estimation. Since the source state at time *t* depends on the state of the former *K* time points, we need to redefine the augmented source state as 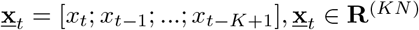 and the augmented dynamic model as Equation 22 to transform VAR(*K*) problem into a VAR(1) one.

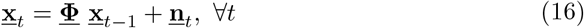

where

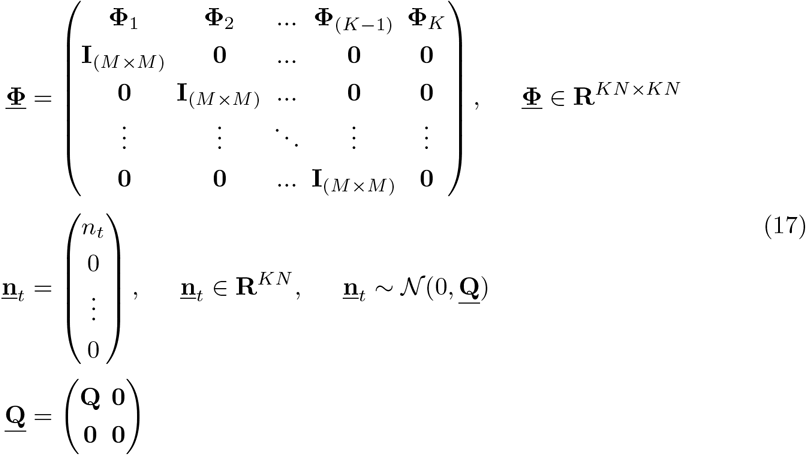

are the augmented state transition matrix and disturbance in source, respectively. And **<u>Q</u>** is the covariance matrix for the augmented disturbance, which is also diagonal, whose diagonal values are the same variance as **Q** and 0 elsewhere. We can also have the augmented version of merged observations as

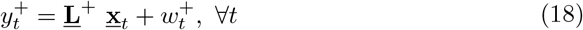

where

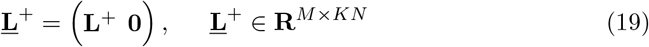

In this way, we can calculate the 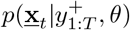 via FIS if the observations as well as *θ* are known.

Next, according to [40] one can define

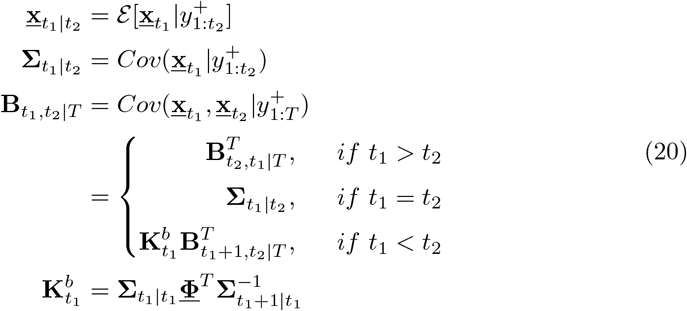

as the mean, covariance matrix as well as the cross-covariance matrix for any given *t*_1_ and *t*_2_. Then we can fit the model with the FIS framework.

In the forward filtering step, we start with the initial value obtained via VAR. Then we can do for *t* = 0, 1, 2, …, *T −* 1

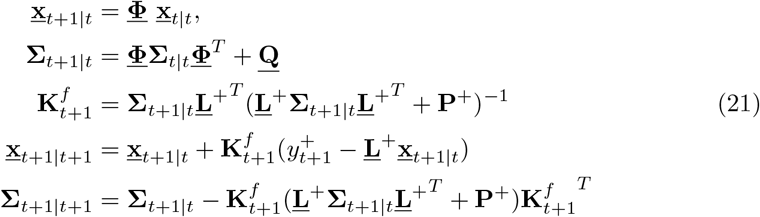

Taking the results from the filtering step, we can further do backward smoothing for

*t* = *T −* 1, *T −* 2, …, 0 as

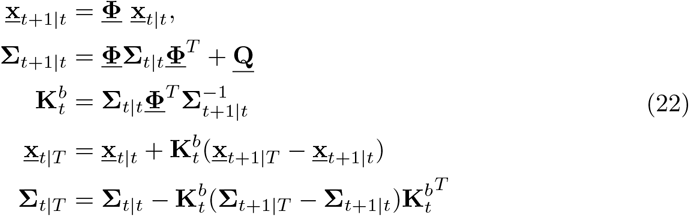

Then we can calculate the cross-covariance matrix according to Equation 20, and finally, simply extract the first *N* rows of 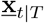and the *N* order submatrix of the upper left corner of the matrix 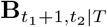 for *∀t*_1_, *t*_2_ = 1, 2, …, *T*, which are exactly the first- and second-order moment of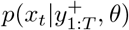 *∀t*, to finalize the E-step in Section 2.2.

#### Algorithm 1

EM Framework for Parameter Estimation

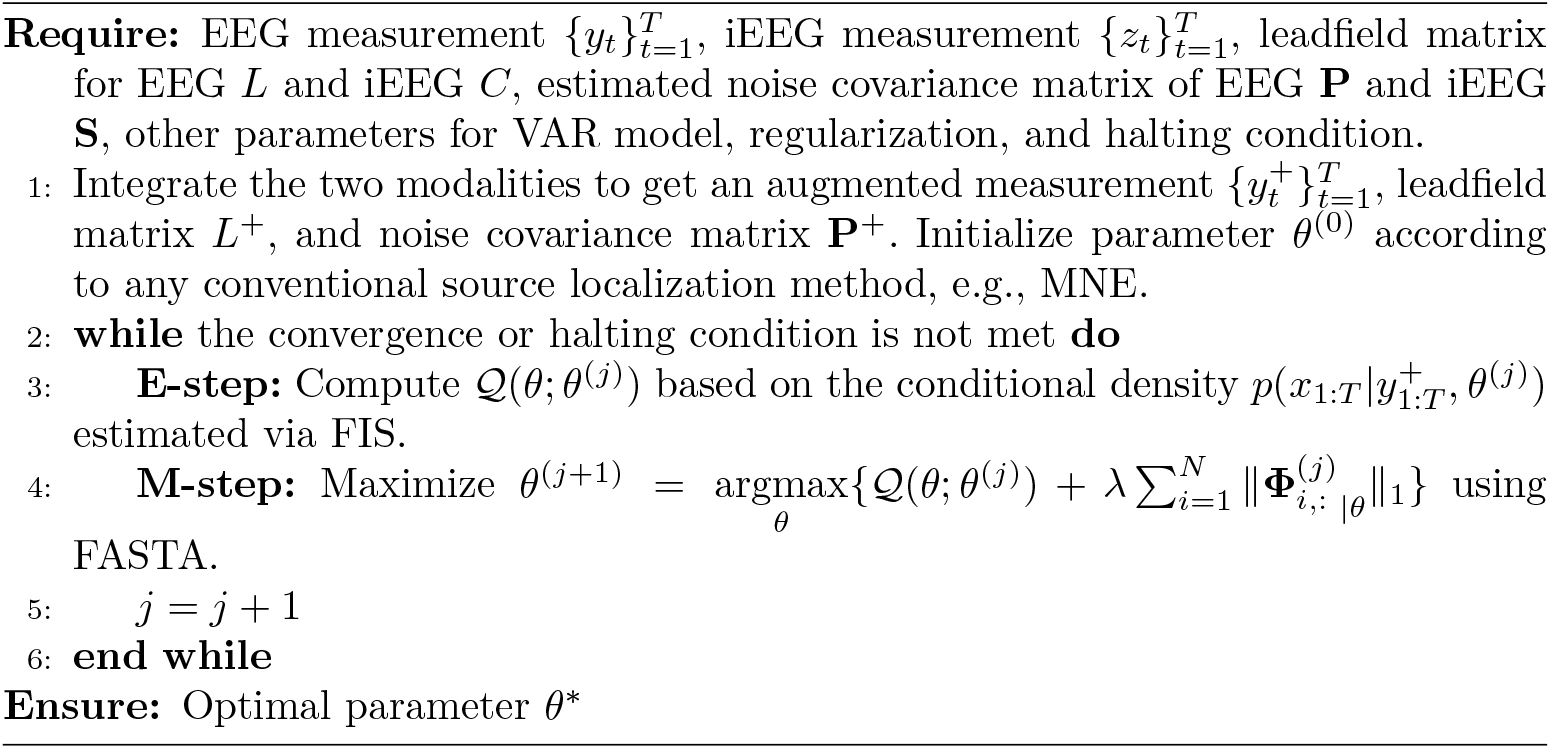

## 3Results

In this study, we first conducted experiments on simulated data and evaluated relevant indicators and results, and then we tested the method on a dataset collected from a real Sternberg verbal working memory task to explore the connectivity maps between cortical and subcortical regions during the encoding and maintenance phases. The experiment setups as well as the evaluation metrics for both simulated and real-data experiments, will be elaborated first, followed by the results and analysis, respectively.

### 3.1 Simulated Experiment

#### 3.1.1 forward model

To generate synthetic EEG data, we used a real head model to compute the leadfield matrix. The T1-MRI images were introduced from FreeSurfer [41] templates. The brain tissue segmentation and source surface reconstruction were conducted using FreeSurfer. Then a three-layer boundary element method (BEM) head was built based on the reconstructed surfaces. A 128-channel BioSemi EEG cap layout was used, and the EEG channels were co-registered with the head model and further validated on the MNE-Python toolbox [42]. The source space was parcelled and clustered according to the ico-1 spacing standard that contains 42 sources in each hemisphere, with 84 sources combined, resulting in a leadfield matrix *L* with a dimension of 128 by 84.

#### 3.1.2 Data Generation and Preprocessing

Next, to generate a simulated time series of a causal series. We referred to the famous Berlin Brain Connectivity Benchmark (BBCB) [43] and slightly simplified it by using the autoregression idea based on *K* randomly generated state transition matrices **Φ** to generate the source signal, and to ensure the source signal does not diverge, each component in **Φ** should be less or equal to 1. Then we add an independent random Gaussian noise to each source signal at every time step. Lastly, an acausal third-order Butterworth filter with zero phase delay was applied with pass bandwidth [0.1Hz, 40Hz].

With the generated source signal, we can easily generate the observations by multiplying the leadfield matrix with the source signal and adding the channel-wise correlated random Gaussian noise according to a given signal-to-noise ratio (SNR). It is worth noting that, if the partial observations are introduced, the leadfield matrix should be augmented as illustrated in 2.3, and based on the assumption that the measurement noise of iEEG should not only be independent of that of the EEG but also have much higher SNR level [44]. Thus the iEEG noise needs to be generated separately under a relatively high SNR, and we can obtain the intact observation noise correlation matrix by then. In this way, all the inputs for the proposed framework are ready.

#### 3.1.3 Evaluation Metrics

In the simulated experiment, the ground truth of the connectivity network is defined based on the generated state transition matrix **Φ**. The easiest way to represent if there is a link between region *i* and *j* is by looking at the summation of *K* **Φ**s and see if the component with index (*i, j*) or/and (*j, i*) is non-zero. To make the problem better interpretive, we set *K* = 1, thus the source signal is similar to a Markov process where the state at time *t* is only dependent on the state at time *t −* 1. In this case, the transition matrix **Φ** itself can also work as the correlation matrix for the source signal. Therefore, if both true network **Φ** and estimated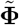have non-zero value on the sample component, we counted it as a good prediction otherwise, a bad one. To evaluate the performance, we defined the sensitivity and accuracy of the connectivity links estimation as

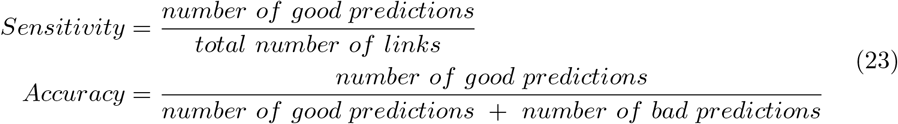

Moreover, the connectivity prediction can be visualized via BrainNet Viewer[45] to provide a more intuitive comparison.

#### 3.1.4 Simulation Setups and Results Analysis

The simulation was conducted in three different manners to thoroughly analyze the effectiveness while introducing iEEG observations in brain network inference. In all experiment setups, the benchmarks were set to the traditional two-step method, which does source localization by taking classical algorithms, such as MNE, sLORETA, etc., first and then applying the Granger causality analysis for each permutation of the estimated source signals. As mentioned before, the evaluation of connectivity is based on the state transition matrix, thus, to further sparsify and robustify the estimated connectivity, we drop all the absolute correlation values that are less than a given threshold, in this simulation, 0.01 was set as the threshold. The observation signals were set with sampling rate 100Hz and a group of 10s time-length sequences with 1000 time steps were generated while the first one second of data were dropped to ensure the stabilization. In order to comprehensively analyze the impact of introducing iEEG, several hyperparameters, i.e., partial observations proportion, SNR, and causal network complexity (which includes the number of activated source regions and the maximum in-degree of each region), were evaluated within a given range. Then we will illustrate each simulation experiment as follows.

##### Impact of Partial Observations Proportion

We first validated how the proportion of partial observations can affect the performance in connectivity estimation. In this case, the independent variable is partial observations proportion, namely the coverage rate, and its range was set from 0% to 50%. The SNR for EEG and iEEG observations were set to −5dB and 30dB, respectively. The network complexity was set with the number of activations as 10 and in-degree as 2. For each parameter combination, 10 repetitive runs were applied to ensure that the results were accurate and reliable. The statistics are shown in Table 1

**Table 1.** Partial Observations Proportion Impact Evaluation.

| Metrics | 0% | 10% | 20% | 30% | 40% | 50% | 60% |
| --- | --- | --- | --- | --- | --- | --- | --- |
| Sensitivity | $0.461 \pm 0.155$ | $0.517 \pm 0.042$ | $0.556 \pm 0.070$ | $0.639 \pm 0.120$ | $0.690 \pm 0.147$ | $0.721 \pm 0.153$ | $0.852 \pm 0.131$ |
| Accuracy | $0.189 \pm 0.176$ | $0.179 \pm 0.161$ | $0.180 \pm 0.174$ | $0.545 \pm 0.224$ | $0.657 \pm 0.194$ | $0.687 \pm 0.184$ | $0.735 \pm 0.201$ |

It is worth noting that as partial observation proportion increases, both sensitivity and accuracy on connectivity estimation show an increasing trend, and the increase, especially for accuracy, is significant at 30%, which proves that when iEEG is introduced, and an appropriate amount of activated area is observed, performance can be significantly improved. A visualization example for 30% partial observation is shown in Figure 2

**Fig. 2.**
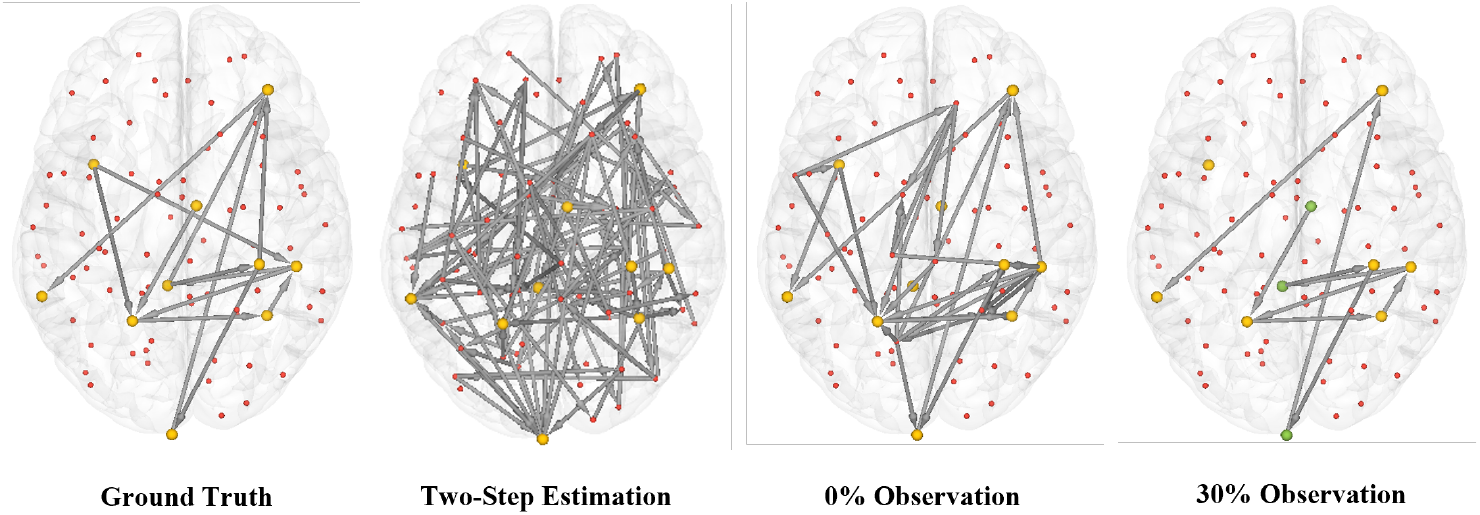
Visualization for network estimation in different methods, where the activated patch centers are highlighted with yellow color, and the iEEG observed patch centered are highlighted with green color. For two-step methods, the eLORETA that has the best performance is taken for comparison.

##### Impact of Observation SNR

The SNR of the observations is another key factor. As mentioned before, here we assume the SNR in the iEEG signal is far greater than that of the EEG signal, plus the iEEG signal should be stable and accurate enough with a relatively high SNR, whereas the EEG signal is much inaccurate and less stable. Therefore, we simply set the iEEG SNR to 30dB as in the previous experiment and changed the SNR for EEG to see how accurate iEEG measurement can improve the estimation performance under different disturbance levels of EEG observations. The SNR for the EEG signal was set from −10dB to 10dB, meanwhile, the iEEG observation proportion was set to 30%, and the network complexity configuration is the same as the previous experiment in that the number of activations is 10 and the in-degree is 2. Table 2 shows the statistical result for this experiment.

**Table 2.**
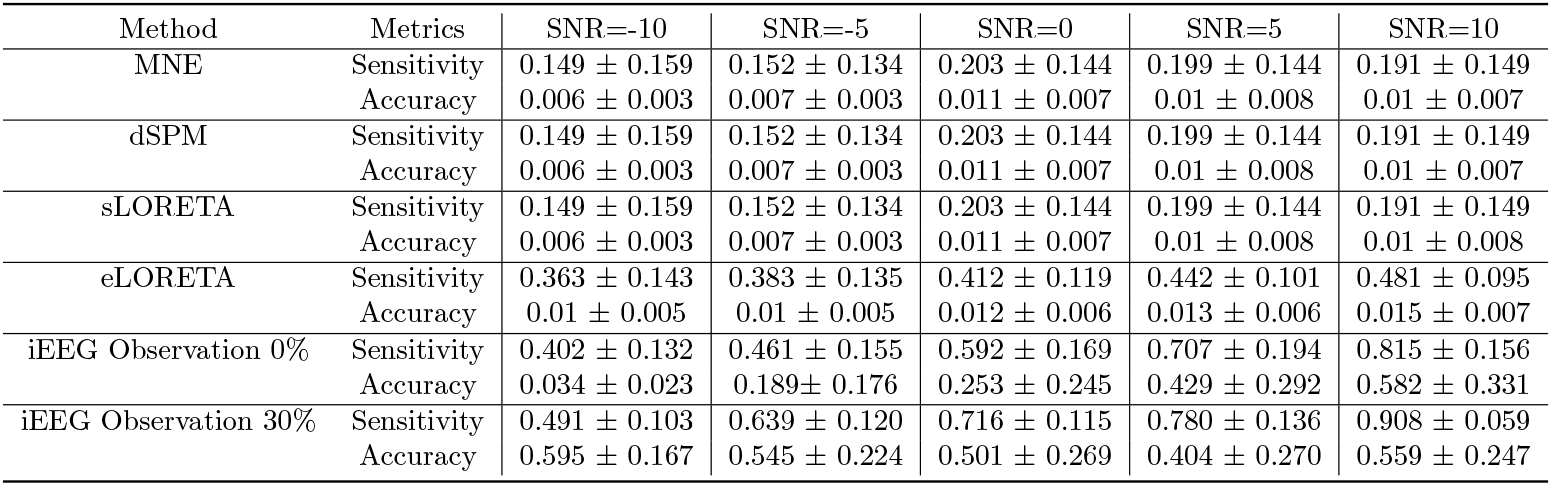
SNR Impact Evaluation.

As we can see from the results, the classical two-step method usually does not perform well when SNR is low, when SNR increases, even if some algorithms, such as eLORETA can capture nearly half of the correct connectivities, the accuracy is still unsatisfying. By contrast, for the dynamic model method when SNR for measurement reaches a relatively high level, the performance of both using and not using iEEG observation has a similar performance. However, when SNR is decreasing, metrics for both cases show a downward trend. But we can find that when SNR is less than zero, the accuracy for no iEEG observation case drops significantly, indicating that in the absence of iEEG observation, a large number of incorrect links in the network are estimated, whereas, with the help of iEEG, the accuracy is relatively stable that is no significant decline in performance, which further prove that by introducing iEEG on can estimate the brain connectivity much accurate, especially under low-quality extracranial measurement circumstances. Therefore, we can conclude that when measurement noise becomes severe, introducing a small proportion of iEEG will not significantly improve correct estimations, i.e. sensitivity, but it can better constrain the incorrect estimated links or, in other words, can achieve a much higher accuracy.

##### Impact of Causal Network Complexity

Lastly, we verified the effect of introducing iEEG observations on the experiment under different causal network complexity circumstances. As mentioned before, the network complexity consists of the activated source number, namely the number of nodes in the network, and the in-degree for each node that affects how many edges are in the network. To mitigate the coupling phenomenon that may influence the results, the two independent variables were changed separately. The number of nodes was changed from 10 to 20, while the in-degree was adjusted from 1 to 3. The other parameters are similar to previous settings in that the iEEG observation proportion was set to 30%, and the SNR for EEG and iEEG observations were set to −5dB and 30dB, respectively.

According to the result, for two-step methods, the performance is usually unsatisfying, and even though the sensitivity is increased while the network complexity increases, they usually suffer from low accuracy or incorrect connectivity prediction. According to our method, we can conclude that whether it is increasing the number of nodes or increasing the number of links to increase the complexity of the network would result in performance degradation. However, due to the existence of iEEG, in most cases, both sensitivity and accuracy, especially the latter, are better than the case where iEEG is missing. This result can be another proof that a more accurate connectivity estimation could be achieved while the iEEG can be collected with EEG simultaneously. The evaluation details can be found in Table 3 and 4.

**Table 3.** Number of Nodes Impact Evaluation.

| Method | Metrics | #Nodes=10 | #Nodes=15 | #Nodes=20 |
| --- | --- | --- | --- | --- |
| MNE | Sensitivity | $0.152 \pm 0.134$ | $0.242 \pm 0.186$ | $0.148 \pm 0.144$ |
| | Accuracy | $0.007 \pm 0.003$ | $0.012 \pm 0.008$ | $0.011 \pm 0.005$ |
| dSPM | Sensitivity | $0.152 \pm 0.134$ | $0.254 \pm 0.195$ | $0.208 \pm 0.212$ |
| | Accuracy | $0.007 \pm 0.003$ | $0.012 \pm 0.009$ | $0.01 \pm 0.005$ |
| sLORETA | Sensitivity | $0.152 \pm 0.134$ | $0.254 \pm 0.195$ | $0.208 \pm 0.212$ |
| | Accuracy | $0.007 \pm 0.003$ | $0.012 \pm 0.009$ | $0.01 \pm 0.005$ |
| eLORETA | Sensitivity | $0.383 \pm 0.135$ | $0.39 \pm 0.169$ | $0.389 \pm 0.146$ |
| | Accuracy | $0.01 \pm 0.005$ | $0.009 \pm 0.005$ | $0.011 \pm 0.005$ |
| iEEG Observation 0% | Sensitivity | $0.461 \pm 0.155$ | $0.350 \pm 0.087$ | $0.398 \pm 0.096$ |
| | Accuracy | $0.189 \pm 0.176$ | $0.118 \pm 0.095$ | $0.191 \pm 0.091$ |
| iEEG Observation 30% | Sensitivity | $0.639 \pm 0.120$ | $0.518 \pm 0.110$ | $0.547 \pm 0.093$ |
| | Accuracy | $0.545 \pm 0.224$ | $0.500 \pm 0.172$ | $0.552 \pm 0.182$ |

**Table 4.** In-degree of Nodes Impact Evaluation.

| Method | Metrics | In-degree=1 | In-degree=2 | In-degree=3 | In-degree=4 |
| --- | --- | --- | --- | --- | --- |
| MNE | Sensitivity | $0.121 \pm 0.037$ | $0.152 \pm 0.134$ | $0.221 \pm 0.19$ | $0.298 \pm 0.271$ |
| | Accuracy | $0.005 \pm 0.002$ | $0.007 \pm 0.003$ | $0.014 \pm 0.012$ | $0.015 \pm 0.011$ |
| dSPM | Sensitivity | $0.121 \pm 0.037$ | $0.152 \pm 0.134$ | $0.423 \pm 0.294$ | $0.504 \pm 0.296$ |
| | Accuracy | $0.005 \pm 0.002$ | $0.007 \pm 0.003$ | $0.011 \pm 0.01$ | $0.011 \pm 0.009$ |
| sLORETA | Sensitivity | $0.121 \pm 0.037$ | $0.152 \pm 0.134$ | $0.423 \pm 0.294$ | $0.504 \pm 0.296$ |
| | Accuracy | $0.005 \pm 0.002$ | $0.007 \pm 0.003$ | $0.011 \pm 0.01$ | $0.011 \pm 0.009$ |
| eLORETA | Sensitivity | $0.196 \pm 0.098$ | $0.383 \pm 0.135$ | $0.505 \pm 0.189$ | $0.63 \pm 0.139$ |
| | Accuracy | $0.005 \pm 0.003$ | $0.01 \pm 0.005$ | $0.01 \pm 0.007$ | $0.009 \pm 0.006$ |
| iEEG Observation 0% | Sensitivity | $0.57 \pm 0.194$ | $0.461 \pm 0.155$ | $0.401 \pm 0.198$ | $0.422 \pm 0.241$ |
| | Accuracy | $0.228 \pm 0.234$ | $0.189 \pm 0.176$ | $0.149 \pm 0.201$ | $0.109 \pm 0.168$ |
| iEEG Observation 30% | Sensitivity | $0.797 \pm 0.129$ | $0.639 \pm 0.120$ | $0.546 \pm 0.143$ | $0.422 \pm 0.203$ |
| | Accuracy | $0.697 \pm 0.248$ | $0.545 \pm 0.224$ | $0.498 \pm 0.159$ | $0.41 \pm 0.215$ |

### 3.2 Real Data Experiment - Cortical-subcortical Information Flow During Verbal Working Memory Task

Working Memory (WM) is commonly associated with learning, understanding, executive functioning, information processing, intelligence, and problem-solving in humans and a variety of animals from infancy to old age [46]. Maintaining content in WM requires communication within an extensive network of brain regions [47]. In this section, we applied the proposed method to estimate and analyze the brain network between cortical and subcortical regions at different WM processes, particularly the encoding and maintenance process, in a verbal WM task for epilepsy patients.

#### 3.2.1 Data Description and Preprocessing

The dataset was collected from 15 subjects during a verbal working memory task, where subjects were epilepsy patients undergoing intracranial monitoring for localization of epileptic seizures. Subjects performed a modified Sternberg task in which the encoding of memory items, maintenance, and recall were temporally separated. The dataset includes simultaneously recorded scalp EEG with the 10-20 system, iEEG recorded with depth electrodes, waveforms, and the MNI coordinates and anatomical labels of all intracranial electrodes [48]. Each subject contains several sessions, while for each session, there are 50 events. Considering the possibility of similar experimental results distribution, for subjects that contain more than 4 sessions, we discard the excess parts to make sure there are enough samples for each subject and reduce the redundant computation. Moreover, there is an 8s recording with 200 Hz and 2000 Hz sampling rates for EEG and iEEG in an event, respectively, where 0-1 s represents the fixation phase, 1-3 s is the encoding phase in WM, 3-6 s records the maintenance phase in WM, and 6-8s is for response phase. In this paper, we only focus on the encoding and maintenance phases, and due to the possibility that the encoding phase could exceed the expected time interval, which is caused by the auditory encoding process that may extend past the visual stimulus in the experiment, only the last 2 s of maintenance should be preserved [47].

Since there is no MRI data for each subject, thus a template head MRI from FreeSurfer was applied. We then used MNE-Python to calculate the forward model based on the ’fsaverage’ template [49] over the oct-5 source space, where each hemisphere resulted in about 2052 patches. Since the electrodes for each subject are different, the forward solution was calculated independently according to the standard 10-20 montage for the electrodes used by each subject. To analyze the connectivity among different functional regions, we introduced the Harvard-Oxford (HO) atlas that contains 48 cortical and 21 subcortical structural areas derived from anatomical structural data and segmentations, and all the source locations from the forward model were aggregated into each atlas area by following the nearest neighbor principle in MNI space. In this way, we transformed the original source space with high dimensions into a lower-dimensional atlas source space. Meanwhile, we averaged the columns in the EEG leadfield matrix based on the source aggregation results for the sources that were merged into the same atlas area. Similar processing steps were applied to iEEG data. The iEEG channels were firstly mapped onto HO atlas areas based on the electrodes’ positions, and measurement signals were averaged in accordance with the channels that were assigned to the same atlas area by then. The row of leadfield matrix for iEEG that represents the mapping from source signal to electrode measurement was set as a unit vector with a value of 1 at the assigned atlas area and 0 otherwise for each electrode. Since there are no empty room recordings for the EEG and iEEG device from the dataset, in this work, we assume the noise of EEG as a multivariate Gaussian with 0 mean and standard deviation 1, and iEEG as a multivariate Gaussian with 0 mean and standard deviation 0.1. Lastly, both EEG and iEEG signals were down-sampled at a 50 Hz sampling rate.

#### 3.2.2 Results

In the experiment, we used an order-2 VAR model for the EM estimation framework mentioned in section 2.2. We estimated the regional connectivity of brain activity during encoding and maintenance phases separately for each human participant’s tasks. Then we took the sum over the estimated state transition matrices for each task and obtained the aggregated result by averaging all task-leveled state transition matrices according to different phases. To ensure the stability of the result, we only keep the connection from region *i* to *j* that contains signal power no less than 10% of the signal power from region *j* to itself, i.e., 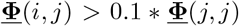. Moreover, we further merged the HO atlas regions in the cortical area into lobe-leveled granularity, while caudate, putamen, and pallidum were aggregated as basal ganglia to pursue robust results further.

The estimated dynamic network for encoding and maintenance phases is shown in Figure. 3. The results showed a significant difference in the network between the two phases. In the encoding phase, the results revealed that there are mainly connections from the cortical to subcortical regions. Specifically, information flows from the frontal, temporal (including regions that are related to auditory processing), and parietal lobes to the thalamus and basal ganglia. On the other hand, the connections that start from the subcortical region mainly point to its internal areas. The results showed a direction from the basal ganglia and hippocampus to the thalamus in the encoding phase. Besides, we can also find there exist connections that start from the amygdala to the basal ganglia and thalamus. By contrast, during the maintenance phase, the main difference lay in the connections originating from subcortical regions compared to the encoding phase. A significant reverse direction from the thalamus and basal ganglia to the frontal and parietal lobes can be found, and there is also a connection from the hippocampus to the temporal lobe (including the auditory processing regions) in this phase.

**Fig. 3.**
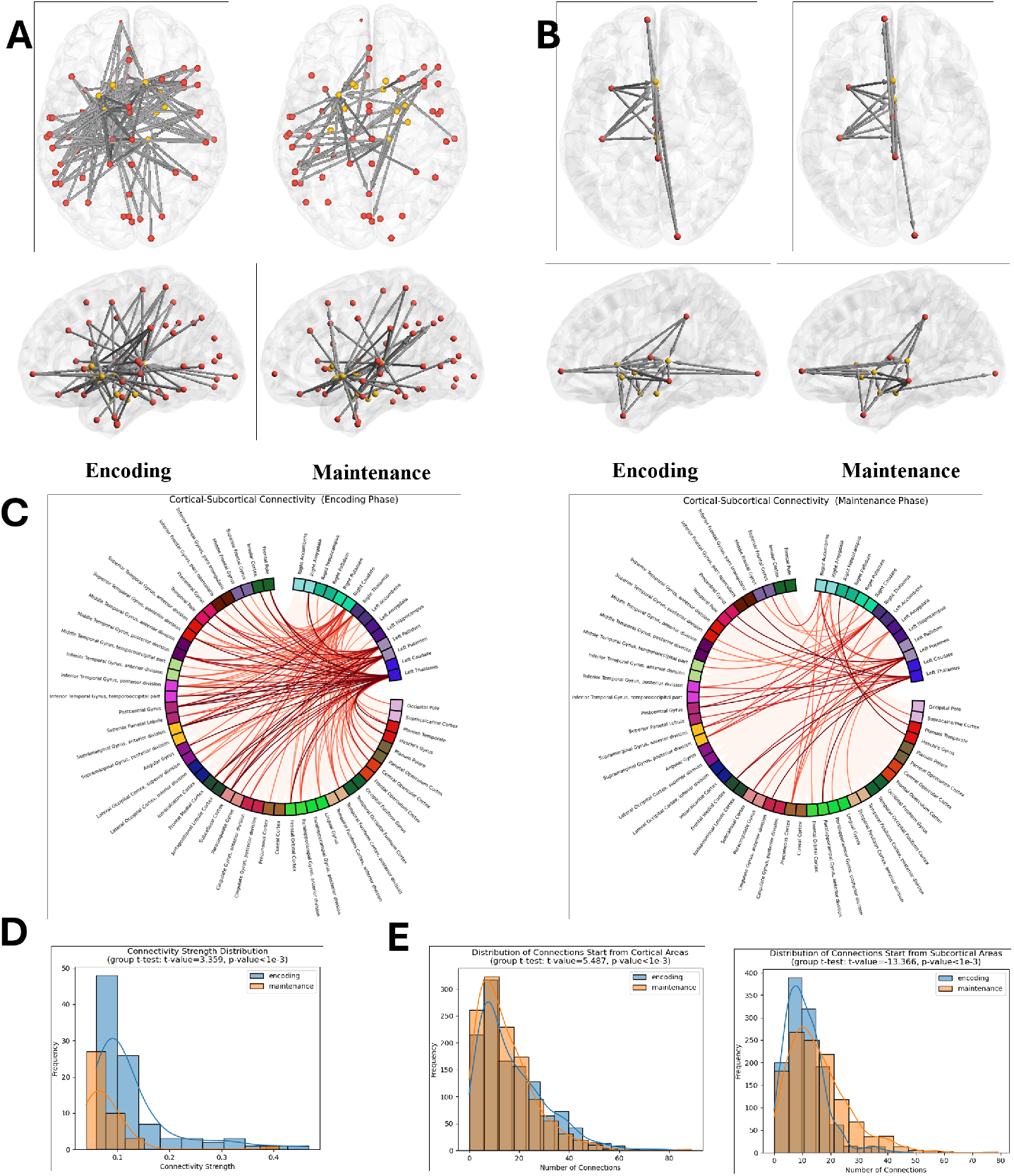
Cortical-subcortical connectivity estimation during encoding and maintenance phases in the verbal working memory task. Sub-figure A shows the brain connectivity network in two WM phases according to the HO atlas, while B is the network of aggregated lobe-leveled regions used for macroscopic expression. C is the circular brain connectivity graph in HO atlas parcellation for encoding and maintenance phases, respectively. D shows the distribution of absolute connectivity strength between cortical and subcortical regions during encoding and maintenance phases based on the averaged results, and a group t-test was applied with a t-value of 3.359 and p *<* 0.001. Lastly, E presents the distribution of connection numbers during two phases according to the event-leveled results, where the figure on the left represents the distribution of the number of the information flows starting from the cortical regions, and a group t-test gave a t-value of 5.487 and p *<* 0.001 between the two phases, whereas the right one presents the information flows start from the subcortical regions and with a t-value of −13.336 and p *<* 0.001 for group t-test.

#### 3.2.3 Discussion

To our knowledge, this is the first inference framework of the whole brain connectivity map during the encoding and maintenance phases of the Sternberg WM task based on simultaneously collected EEG and iEEG. Our study not only helps to infer the brain-wide connectivity network for the WM process, but it can also be applied to the research of other similar problems. It provides new insight into the importance of a clinically integrated collection of iEEG and EEG information, as well as an in-depth exploration of the integration modeling and inference framework

The results of this study showed the activation and dynamic network of cortical and subcortical regions during a verbal working memory task. The results not only showed the same cortical regions activated in the verbal WM task as in traditional studies, such as frontal regions, but also presented subcortical-related connections, which have been demonstrated in previous related studies.

Firstly, a similar result was found in the information flow between cortical and subcortical areas during the encoding and maintenance phases that related to an auditory encoding process in the WM task as in the previous study, which showed the information flows from the temporal lobe to the hippocampus in the encoding phase. A reverse direction appears in the maintenance phase [47]. Although, in our estimation, there is no significant result revealing the connection from the temporal lobe to the hippocampus in the encoding phase, which is likely caused by the position deviation during atlas region aggregation in the preprocessing stage, we still find the information flow from the temporal lobe to other memory processing regions that are adjacent to hippocampus, e.g., caudate, which has also been proved to have synergy with hippocampus in the WM process [50]. While in the maintenance phase, the same reverse information flow from the hippocampus to the temporal lobe was shown in our estimation.

Since it is generally believed that WM is resource-limited, therefore, it is reasonable that the brain would strive to enhance WM in order to facilitate attention, temporary storage, and manipulation of target material [51, 52]. The results obtained by our method are consistent with previous studies, both indicating that the basal ganglia in verbal WM play a role in enhancing goals and inhibiting distracting factors, especially the caudate part is continuously activated during the encoding stage.

Previous research employing fMRI only to investigate language generation in healthy individuals has revealed not only activity in the right caudate nucleus and putamen but also activation in related regions within the left hemisphere [53]. These findings suggested that left basal ganglia activity plays a role in augmenting language processing in the dominant hemisphere, while right basal ganglia activity serves to suppress potentially disruptive activation in the non-dominant right frontal cortex. The same phenomenon is presented in our results, which suggest that bilateral basal ganglia is involved in activities during different phases of WM.

Besides, previous work also proved that not only the basal ganglia but also the thalamus is active during the encoding and maintenance phases of a verbal WM task [51], and our result was consistent with the previous study, where the basal ganglia, especially the caudate, and thalamus are active during the encoding and maintenance phase, and we found there is an information flow from frontal, temporal and parietal lobes to the thalamus and basal ganglia in the encoding phase and a reverse directional flow in the maintenance phase.

Moreover, previous studies have suggested lots of evidence of the amygdala contributing to memory processing, for it plays an important role in the state-dependent encoding of exploratory behavior [54], and contains a higher proportion of response-specific concept cells compared to the hippocampus [55]. We also found that the amygdala is active in our results, particularly during the encoding process, which is similar to the previous work documented that the amygdala was highly activated in the verbal WM task [56].

We further analyzed the connectivity strength and the direction of information flow between cortical and subcortical regions during the encoding and maintenance phases. As shown in sub-figures D and E of Figure 3, we found that the overall connectivity strength during the encoding phase is stronger than that of the maintenance phase. Besides, the amount of connection that flows start from cortical regions is generally greater during the encoding phase than during the maintenance phase, whereas brain connections during the maintenance phase are more active in the information flow that is directed from subcortical regions to elsewhere. A set of group t-tests was applied as proof for the difference significance of the findings as stated in the annotation of Figure 3.

## Declarations

The authors declare no competing interests.

## 4 Conclusion

In this study, we proposed a method to model the dynamic network of the brain by integrating simultaneously measured scalp EEG and iEEG, which is solved based on the EM framework. Unlike many previous brain network estimation approaches based on fMRI, this method only uses scalp EEG and iEEG, which provide the possibility for real-time sampling and analysis in clinical settings. The proposed methodology enjoys the advantage of information complementary between two modalities. With the appropriate deployment of iEEG, even if the EEG signal has a low SNR level, one can still reconstruct the brain connectivity map with higher accuracy and sensitivity compared to previous works that used scalp EEG or iEEG signal only. Numerical experiments demonstrated that the proposed method performs well under high network complexity cases and yields good robustness with low SNR levels. on a real Sternberg verbal working memory experimental data, the proposed method not only obtained similar results to other studies using the same data cohort but also had strong consistency with the findings of previous related studies. Therefore, this work provides insights into the simultaneous acquisition of EEG and iEEG signals in future research related to brain network analysis and clinical lesion localization.

